# Probing the effects of nonannular lipid binding on the stability of the calcium pump SERCA

**DOI:** 10.1101/470948

**Authors:** L. Michel Espinoza-Fonseca

**Affiliations:** Center for Arrhythmia Research, Department of Internal Medicine, Division of Cardiovascular Medicine, University of Michigan, Ann Arbor, MI 48109, USA

## Abstract

The calcium pump SERCA is a transmembrane protein that is critical for calcium transport in cells. SERCA resides in an environment made up largely by the lipid bilayer, so lipids play a central role on its stability and function. Studies have provided insights into the effects of annular and bulk lipids on SERCA activation, but the role of a nonannular lipid site in the E2 intermediate state remains elusive. Here, we have performed microsecond molecular dynamics (MD) simulations to probe the effects of nonannular lipid binding on the stability and structural dynamics of the E2 state of SERCA. We found that the structural integrity and stability of the E2 state is independent of nonannular lipid binding, and that occupancy of a lipid molecule at this site does not modulate destabilization of the E2 state, a step required to initiate the transition toward the competent E1 state. We also found that binding of the nonannular lipid does not induce direct allosteric control of the intrinsic functional dynamics the E2 state. We conclude that nonannular lipid binding is not necessary for the stability of the E2 state, but we speculate that it becomes functionally significant during the E2-to-E1 transition of the pump.

## Introduction

The Ca^2+^-ATPase SERCA is an ATP-powered transmembrane pump that resides in the endoplasmic reticulum (ER) or in the sarcoplasmic reticulum (SR) of eukaryotic cells. SERCA is one of the most important contributors to Ca^2+^ mobilization across the ER/SR, thus playing a dominant role in Ca^2+^ homeostasis and muscle contractility^1^. In each cycle of its operation, SERCA utilizes the energy produced by hydrolysis of ATP to transport two Ca^2+^ ions into the ER/SR lumen^2,3^. At physiological conditions, SERCA primarily populates a high Ca^2+^-affinity structural state of the pump, E1^4^. As Ca^2+^ concentrations in the cytosol increase, this E1 state binds two Ca^2+^ ions bind to the transport sites, thus inducing ATP hydrolysis and phosphorylation of residue Asp351^5^. These events induce a structural transition toward a phosphorylated, low Ca^2+^-affinity structural state of the pump, E2-P, thus coupling ATP hydrolysis with Ca^2+^ release into the ER/SR lumen^6,7^. Upon Ca^2+^ release, SERCA becomes dephosphorylated and binds 1-2 protons from the ER/SR lumen to neutralize the highly charged transport site and preserve the structural stability of the pump^6^. SERCA in the E2 state releases 1-2 protons from the transport sites to the cytosol, thus completing a cycle of Ca^2+^ and H^+^ countertransport across the ER/SR. Finally, proton-metal ion exchange destabilizes E2 and accelerates the structural transitions toward E1 state required for the next Ca^2+^ pumping cycle^5,7-10^. A scheme of the transport cycle of SERCA is shown in Figure 1A.

**Figure 1.**
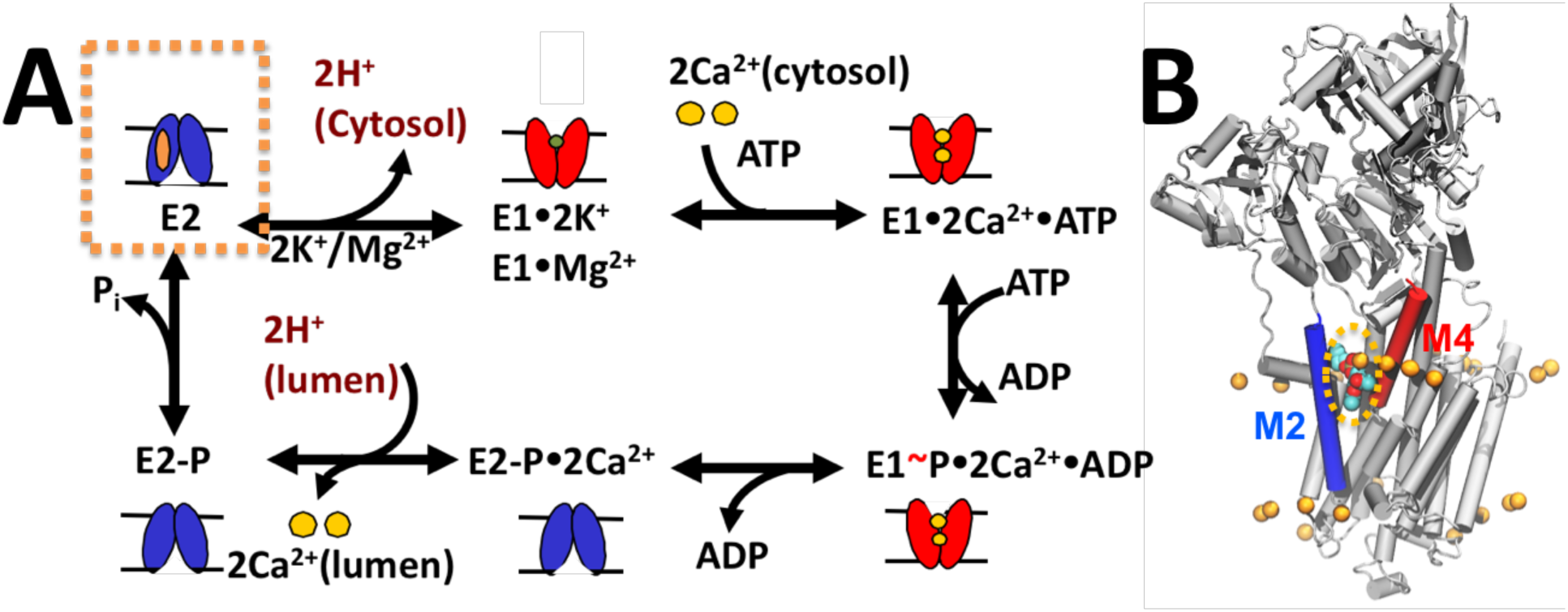
Schematic representation of the functional cycle of SERCA. (A) The scheme shows major low-Ca^2+^ affinity (E2, blue) and high-Ca^2+^ affinity (E1, red) biochemical intermediates of SERCA that are populated during the catalytic cycle of the pump. The orange box shows the E2 state, the only structural intermediate known to bind a nonannular. (B) Structure of the nonannular lipid-bound E2 state of SERCA (gray cartoon). Here, the nonannular lipid, shown as spheres, binds at the interface between transmembrane helices M2 (blue) and M4 (red). The orange spheres show the location of the annular lipid headgroups that surround the transmembrane helices of SERCA; the location of the annular lipids was determined by x-ray crystallography^34^.

The complex mechanism for Ca^2+^ transport by SERCA and the transition from one intermediate to another is influenced by metal ions^11-15^, nucleotide binding^16^, protonation/deprotonation of the transport sites^10,17^, posttranslational modifications^18,19^, and endogenous regulatory proteins^20-25^. However, SERCA also operates in an environment made up largely by the lipid bilayer, so lipids and cholesterol play a central role on SERCA activity through their diverse chemical spectrum and the cooperative physical properties of lipid mixtures of variable composition. For example, spectroscopy experiments have shown that membrane thickness has a direct effect on maximal velocity (V_max_) of the pump, composition of the headgroup induces changes in Ca^2+^ affinity as well as regulation by phospholamban (PLB) in the SR, and that lipid saturation influence both V_max_ and Ca^2+^ affinity^26,27^. Experiments on mice have shown that a change in the balance of the phosphatidylcholine and phosphatidylethanolamine composition in liver ER impairs the Ca^2+^ transport activity of SERCA, indicating that changes in the lipid composition balance can significantly perturb SERCA function in the cell ^28^. More recently, it was shown that changes in the saturated fatty acid on protect SERCA from thermal inactivation, thus showing that annular lipids determine SERCA’s susceptibility to oxidative damage^29^. Cholesterol, which is present in the ER membrane at low concentrations, has also been shown to affect SERCA activity^30^.

These studies have provided insights into the effects of annular and bulk lipids on the activity of SERCA, but the role of nonannular lipids remains elusive. Nonannular bind to distinct hydrophobic sites of membrane proteins or membrane protein complexes, and fulfill a diverse range of functions, from structural stability to allosteric effectors^31,32^. SERCA-lipid interactions have been extensively mapped along the catalytic cycle of the pump^33-36^. These studies revealed the existence of a nonannular lipid binding site located in a cavity formed between transmembrane helices M2 and M4 on the cytosolic leaflet of the ER/SR membrane (Figure 1B)^11,35-38^. More than 50 structures of SERCA representing different structural and biochemical intermediates along the Ca^2+^ pumping cycle have been solved using x-ray crystallography, and this vast amount of structural information has shown that the nonannular lipid binding site is exclusively populated in the protonated, unphosphorylated E2 state of the pump^34,35^(Figure 1). Previous x-ray crystallography studies have suggested that a single nonannular lipid binds to the E2 state and acts as a wedge to keep M2 and M4 apart, thus stabilizing the E2 state of the pump^35^.

While these studies have demonstrated the existence of a nonannular lipid site in the transmembrane domain of SERCA, there is limited information regarding the specific function of nonannular lipids on SERCA stability and function. In this study, we used atomistic molecular simulations to determine the functional outcomes of nonannular lipid binding on the E2 state of the pump. First, we used a crystal structure of the E2 state SERCA as a starting structure to obtain a model of the full length nonannular lipid bound to the transmembrane domain of the pump. We used this structural model to perform all-atom, microsecond molecular dynamics (MD) simulations of the nonannular-SERCA complex at physiological conditions. We also performed a complementary MD simulation of the E2 state in the absence of nonannular lipid. These long-scale simulations were used to systematically probe the effects of non the effects of nonannular lipid binding on the stability and structural dynamics of the protonated E2 state of SERCA.

## Results

### Nonannular SERCA-lipid interactions

We first analyzed the behavior of a single 1-palmitoyl-2-oleoyl-*sn*-glycero-3-phosphocholine (POPC) lipid molecule bound to the nonannular site of SERCA at physiological conditions. We found that the lipid molecule remains bound to SERCA for the entire simulation time, and that there is no lipid exchange between the nonannular site and annulus in the time scales studied here. Clustering analysis showed that the hydrophilic headgroup populates an orientation similar to that resolved by x-ray crystallography (Figure 2A), but the headgroup departs from the initial position to populate different spatial orientations in the microsecond timescale. In this case, the lipid headgroup adopts three main orientations: (i) an orientation in which the headgroup roughly overlaps the position of the crystallographic lipid along the normal axis, but is deep inserted into the nonannular site (Figure 2B); (ii) a low-sitting position of the lipid, with the glycerol group protruding into the annulus (Figure 2C); and (iii) an infrequent (<8%) position of the lipid where the headgroup is shifted by at least 0.5 nm along the z-axis, resulting in the lipid moving closer to the cytosolic domain of the pump (Figure 2D). Previous crystallographic studies have shown that the electron density at the nonannular site is best defined in the presence of 18:1 phosphatidylcholine (DOPC), and that shorter or longer lipids are less ordered when bound to the nonannular site^36^. These experimental evidence helps explain the nonannular lipid mobility observed in our simulation, as the shorter palmitoyl chain (16:0) in the *sn-*1 position of POPC is probably a contributing factor to the mobility of the lipid in this site.

**Figure 2.**
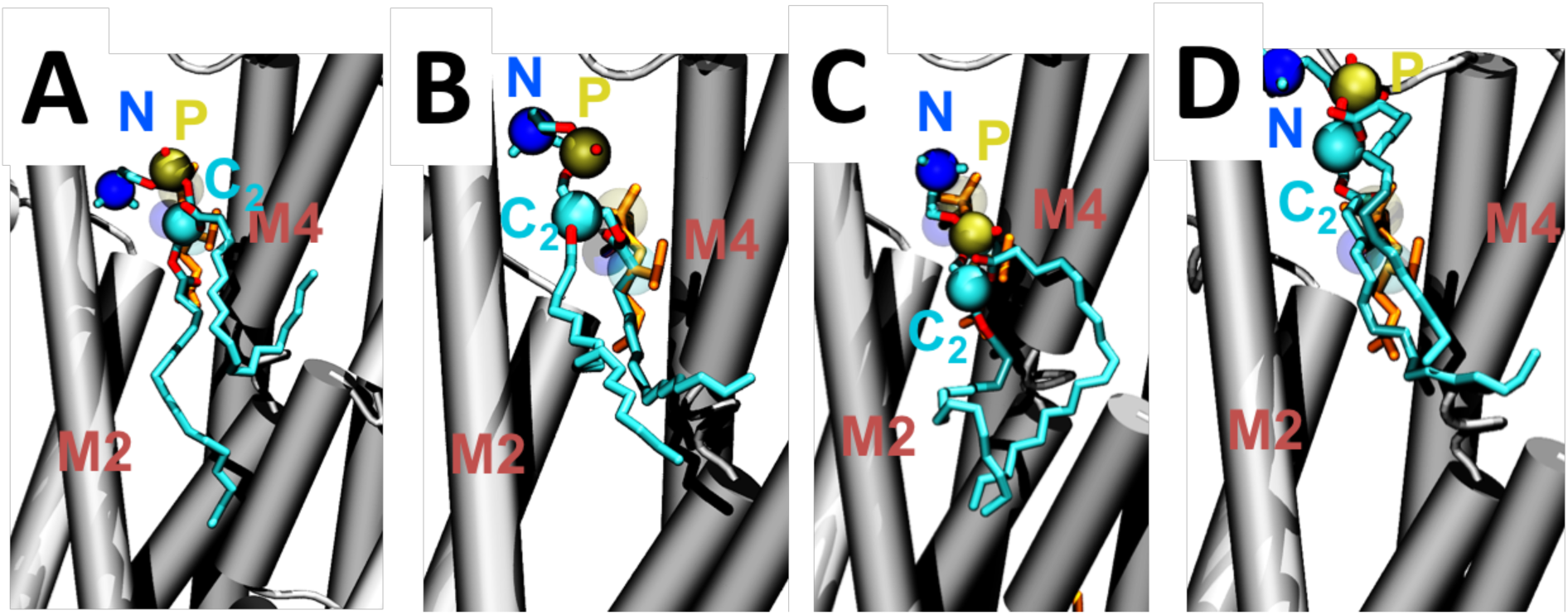
Orientations adopted by the POPC lipid in the nonannular site of SERCA. (A) In this configuration, the headgroup of the lipid roughly overlaps the position of that in the crystal structure of the pump^35^; (B) the axial position of the lipid in this position is similar to ‘A’, but the headgroup is reaches deeper into the nonannular site; (C) In this orientation, the lipid is found in a low-sitting position, where the glycerol group protruding into the annulus; (D) the lipid binds to a position where the headgroup is located closer to the cytosolic domain of the pump. In all panels, SERCA is shown as cartoons, the nonannular lipid as sticks, and the nitrogen, phosphate and carbon-2 atoms of the lipid as spheres.

Clustering analysis indicates that the polar headgroup of POPC is mobile at physiological conditions, and this lipid is not bound in a specific orientation to the nonannular site of the pump. This suggests that there are no specific intermolecular contacts that serve as an anchor for tight binding of the lipid molecule onto the nonannular site. Superposition of reported x-ray crystal structures of complexes of the SERCA-nonannular lipid in the presence [Protein Data Bank (PDB) entries 2agv^35^ and 5xab^34^] and absence (PDB entry 3w5c^11^) of exogenous inhibitors revealed that there are no basic residues vicinity of the lipid (e.g., arginine or lysine) that help stabilization of the complex via salt-bridge interactions. Instead, the position of the nonannular lipid molecule appears stabilized primarily by interactions between residue Gln108 and the phosphate group of the lipid, and by interactions between residues Asn101 or Gln56 and the headgroup of the lipid (Figure 3A). Therefore, we measured the distances between these SERCA residues and the phosphate or choline group of the POPC lipid to determine if these intermolecular interactions are preserved at physiological-like conditions and in the microsecond timescale.

**Figure 3.**
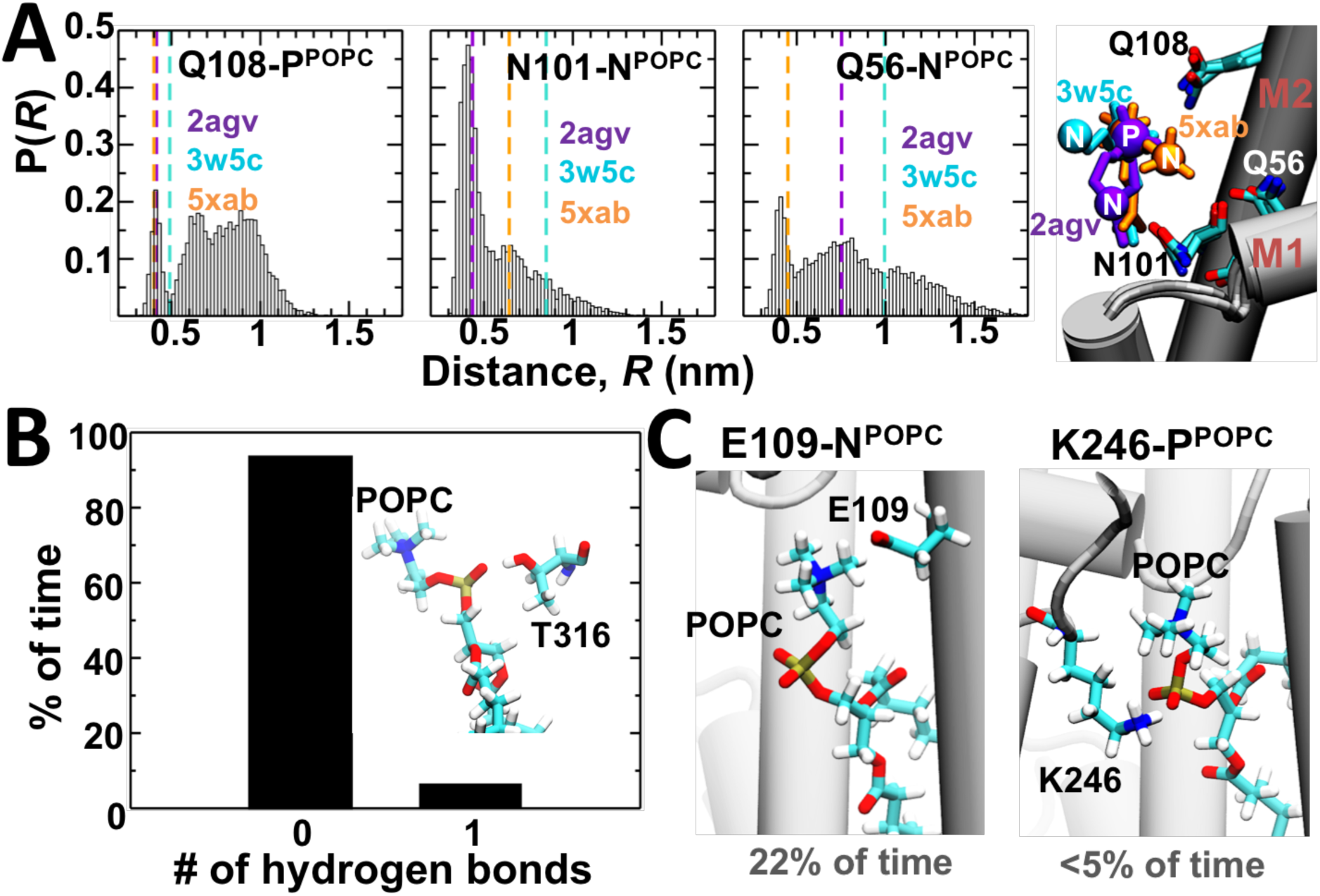
Intermolecular interactions between SERCA and the nonannular lipid. (A) Distance probability distributions for the amide group of residues Gln108, Asn101 and Gln56 of SERCA and the nitrogen or phosphate atoms of the nonannular lipid. The dashed lines indicate the distances calculated from representative crystal structures of the E2 state (right panel). (B) Stability of the hydrogen bond formed between the phosphate group of the nonannular lipid and residue Thr316 of SERCA. For this intermolecular interaction, the presence and absence of a single hydrogen bond is expressed as a percentage of time. (C) Formation of electrostatically favorable contacts between residues Glu109 and Lys246 of SERCA and the nonannular lipid; while these intermolecular interactions are not observed in crystal structures, they are present in the simulation at physiological conditions.

Analysis of the SERCA-nonannular lipid distance histograms that all protein-lipid atom pairs yield broad distributions in the microsecond time scale. In the case of the distance between Gln108 and the phosphate group of the lipid, the distance range defined by the crystal structures of SERCA is very narrow (0.4 nm ≤ *R* ≤0.5 nm); here, we found that ~15% of the total intermolecular distances calculated in the MD trajectory of the nonannular lipid-bound SERCA either fall within the boundaries set by the crystal structures of SERCA to form direct interactions between the phosphate group of the lipid and the amine group of Gln108 (*R*≤0.5 nm) (Figure 3A). In contrast, the distance range estimated for the headgroup nitrogen atom and residues Gln56 or Asn101 in the crystal structures is much broader, with distances ≤1 nm and <0.85 nm for the Gln56-N^POPC^ and Asn101-N POPC pairs, respectively (Figure 3A). Comparison with experimental data showed that that 71% of the choline-Gln56 distances and 89% of the choline-Asn101 distances calculated in the MD trajectory fall within the boundaries set by the crystal structures of the E2 state (Figure 3A). Nonetheless, only ~24% of the choline-Gln56 distances and ~60% of the choline-Asn101 distances are within the typical range required for favorable carbonyl-choline interactions (*R*≤0.55 nm)^39^. X-ray crystallography studies also identified the side chain of SERCA residue Thr316 (transmembrane helix M4) as a potential hydrogen bond donor to stabilize the position of the phosphate group in the nonannular site^35^. However, analysis of the trajectory revealed that the hydrogen bond between Thr316 and the phosphate moiety of the nonannular lipid is present only in 6.4% of the simulation time (Figure 3B). We also detected the formation of favorable contacts between SERCA and the nonannular lipid interactions which are not observed in crystal structures, such as the transient (<5%) interaction between residue Lys246 and the phosphate group of the lipid, and the interaction between residue Glu109 and the choline moiety of the POPC lipid that is formed only in 22% of the structures sampled in the simulation (Figure 3C).

### Structural changes of the helix–helix interface in response to nonannular lipid binding

We determined the changes in the M2-M4 interface in the presence or absence of nonannular lipid. Visualization of the MD trajectories revealed that helices M2 and M4 remain physically separated from each other in the presence of the non-annular lipid (Figure 4A). We found that the in the absence of nonannular lipid, the space separating helices M2 and M4 becomes narrow rapidly in the simulation (t<0.05 μs), thus allowing direct favorable interaction between polar residues from both helices (Figure 4A). We found that upon narrowing of the nonannular site, the headgroups of annular lipids interact with the side chain of residues that might be important for binding (e.g., Gln108). Nevertheless, we did not observe lipid binding to the narrow M2-M4 interface at the time scales covered in this study, thus allowing us to determine changes in the structure of the interface in response to binding and removal of the nonannular lipid.

**Figure 4.**
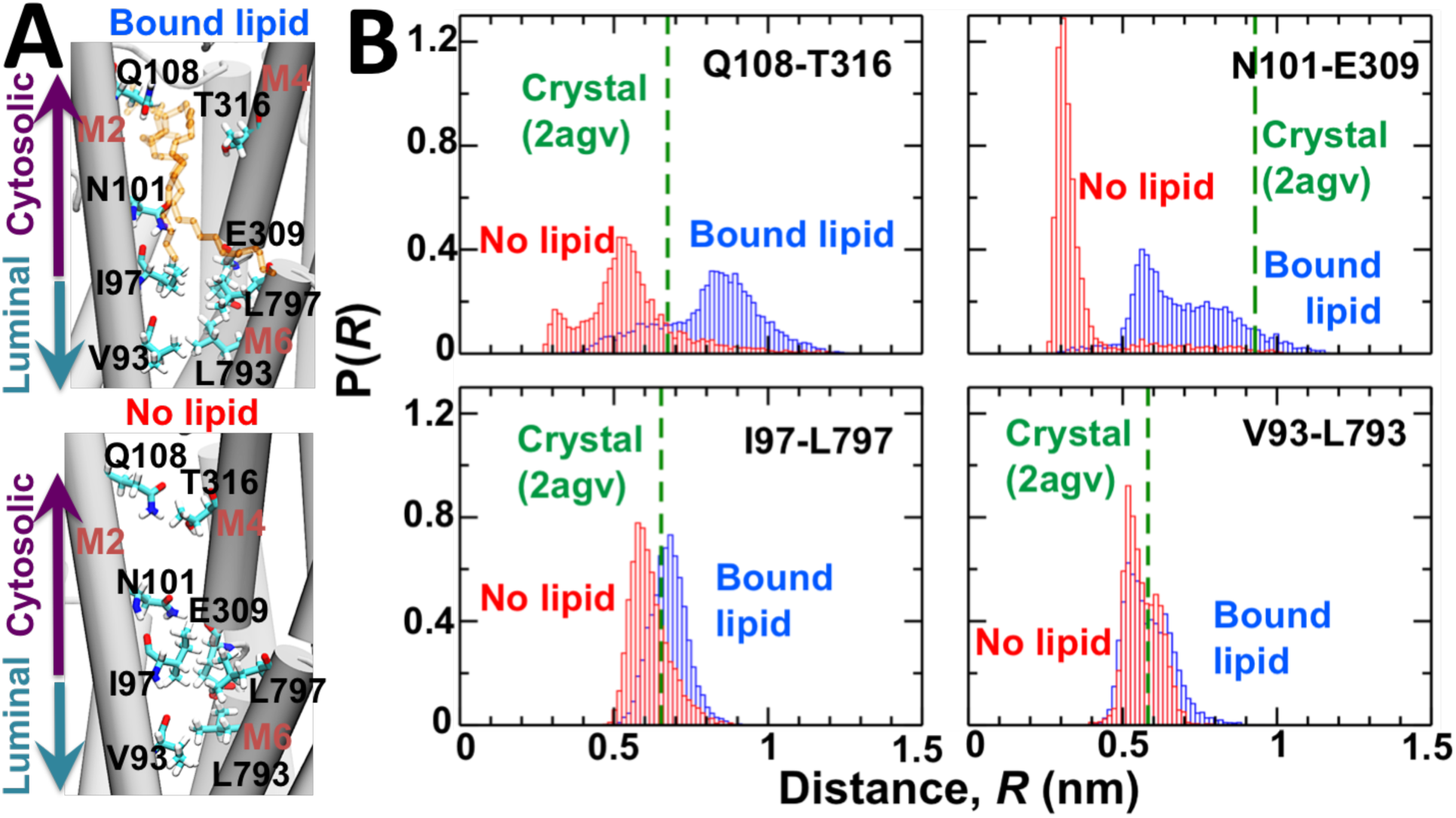
Structural changes of the M2-M4 and M2-M6 interfaces in response to nonannular lipid binding. (A) Structure of transmembrane helices M2, M4 and M6 in the presence (top) and absence (bottom) of nonannular lipid. SERCA is shown as a cartoon, and residues at the helix–helix interface as well as the nonannular lipid are shown as sticks. The location of the cytosolic and luminal sides of the transmembrane domain are shown for clarity. (B) Distance probability distributions for residue pairs Gln108^M2^-Thr316^M4^, Asn101^M2^-Glu309^M4^, Ile97^M2^-Leu797^M6^ and Val93^M2^-Leu793^M6^. Distances were calculated in the presence (blue) and absence (red) of nonannular lipid. The dashed line shows the distance calculated for each residue pair from the crystal structure of the pump^35^.

We measured distance distributions between interhelical residue pairs Gln108^M2^–Thr316^M4^ and Asn101^M2^–Glu309^M4^ to determine the effects of nonannular lipid on the interface between transmembrane helices M2 and M4. We found that the largest changes in the interresidue distance distributions are found in the outermost region of the cytosolic side of SERCA’s transmembrane domain that houses the nonannular lipid site. The histogram plots revealed that in the presence of the nonannular lipid, the cytosolic side of the M2-M4 is mobile, as shown by the broad distance distribution between Gln108^M2^ and Thr316^M4^ (*R*=0.4-1.2 nm); nevertheless, >60% of the interresidue distances are longer than that calculated from the crystal structure of SERCA (*R*=0.65 nm, Figure 4B). In the absence of nonannular lipid, we the interresidue distance Gln108^M2^–Thr316^M4^ also populates a broad distribution in the microsecond time scale; however, we found that >90% of the Gln108^M2^–Thr316^M4^ distances are shorter than that calculated from the crystal structure of SERCA (*R*=0.65 nm, Figure 4B). A substantial lipid binding-induced shift occurs in the distance distribution between residues Asn101^M2^ and Glu309^M4^, which are located near the center of the membrane bilayer (z≈0 nm). Here, the distance histogram for residue pair Asn101^M2^–Glu309^M4^ is characterized by a broad distribution with two well-defined peaks at *R*≈0.5 nm and *R*≈0.8 nm in the presence of nonannular lipid (Figure 4B). However, in the absence of nonannular lipid, there is a redistribution of distances between Asn101^M2^ and Glu309^M4^ toward a sharp, narrow peak at *~*0.3 nm (Figure 4B). In both the presence and absence of nonannular lipid, the distances between Asn101^M2^ and Glu309^M4^ are generally shorter than that calculated from the crystal structure of SERCA, which suggests that this region of the nonannular lipid populates a relatively more compact structure in solution. We also asked whether nonannular lipid binding affects the intermolecular interactions between transmembrane helices M2 and M6, which are located on the luminal side of the nonannular site (Figure 4A). We measured the intramolecular residue pair distances between Ile97^M2^ and Leu797^M6^, and between Val93^M2^ and Leu793^M6^. The distance distribution plots indicate that neither binding nor removal of the nonannular lipid induce changes in the intermolecular interactions between the luminal side of helices M2 and M6 (Figure 4B).

### Effects of nonannular lipid binding on the structural stability and dynamics of SERCA

Our findings show that binding of nonannular lipid induces a spatial separation of the cytosolic regions of helices M2 and M4, thus supporting the hypothesis that this lipid acts as a wedge that keeps apart the cytosolic regions of transmembrane helices M2 and M4^35^. Recent studies have shown that the structural stability of the E2 state is tightly coupled to structural changes in the transmembrane domain, so it is possible that the nonannular lipid-induced separation of helices M2 and M4 is plays a role in maintaining the overall structural stability of the E2 state. To test this hypothesis, we quantitatively compared the architecture of the transmembrane and cytosolic domains of nonannular lipid-bound SERCA with that of the pump in the absence of the nonannular lipid.

We measured the average root mean square deviation (RMSD) for each transmembrane helix of SERCA to determine structural changes in the structure of the transmembrane domain in response to nonannular lipid binding and removal. We found that except for helix M1, the average RMSD in the presence and absence of nonannular lipid are small (<0.2 nm) and similar in magnitude (Figure 5A). Superposition of the structures of SERCA shows that the architecture of nine of the ten transmembrane helices is virtually identical for nonannular lipid-bound SERCA versus nonannular lipid-free SERCA (Figure 5B). We found that helix M1 is substantially more mobile in the lipid-bound protein than in the lipid-free one (Figure 5A), which suggests that binding of the lipid induces changes in the structure of this region of SERCA. Residue Gln56 of SERCA, which is located in the cytosolic region of helix M1, forms favorable interactions with the choline group of the nonannular lipid (Figure 3A), so it is possible that lipid binding is responsible for the structural changes in helix M1 detected in our simulation. However, we found that the cytosolic segment of M1 has low mobility in both trajectories (RMSD<0.1 nm), and that the largest structural shift occurs in the luminal region of helix M1 (Figure 5B). Furthermore, there is a poor correlation between the formation of Gln56-choline interaction and the RMSD of helix M1 (*r*=0.35), which suggests that lipid binding does not have a direct effect on the structural dynamics of helix M1. Extensive (total cumulative time >30 μs) simulations previously reported by our group have shown that the mobility of luminal segment of helix M1 is reproducible, and it occurs regardless of the biochemical state (e.g., bound to metal ions)^13,14^, structural state (e.g., E1 vs E2)^40,41^, or in the presence of regulatory proteins.^40,42^ Therefore, the mobility of M1 observed in the non-annular lipid bound trajectory is most likely due to the inherent flexibility of this helix in solution.

**Figure 5.**
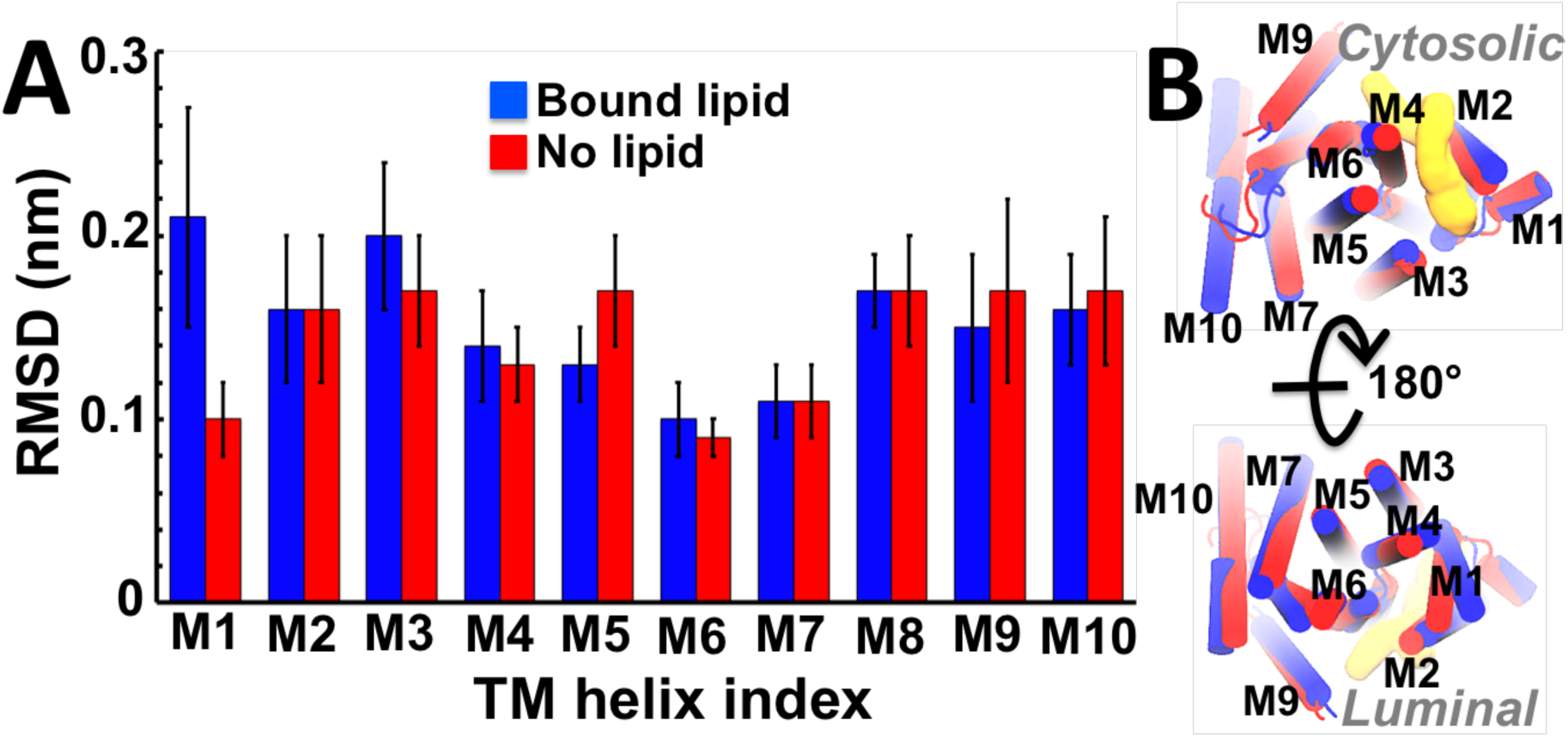
Comparative analysis of the transmembrane helices of SERCA in the presence and absence of nonannular lipid. (A) The RSMD for each TM helix in the presence (blue) and absence (red) of the nonannular lipid was calculated relative to TM helices of the crystal structure of the pump^35^; error bars often represent one standard deviation of uncertainty. (B) Superimposition of the most representative structures of the transmembrane domains of SERCA in the presence (blue cartoon) and absence (red cartoon) of nonannular lipid. For comparison, we show the structure of the cytosolic (top) and luminal (bottom) sides of the transmembrane domain. The location of the nonannular lipid is shown as a yellow surface representation.

Time-dependent RMSD measurements show that the cytosolic domain of SERCA undergoes rapid equilibration (t<0.5 μs) in the MD trajectories, and that the RMSD undergoes moderate fluctuations (RMSD<0.2 nm) in both the presence and absence of nonannular lipid (Figure 6A). Similar to the transmembrane domain, we found overall architecture of this domain is very similar in the presence and absence of nonannular lipid (Figure 6B), which indicates that the nonannular lipid binding is not required for maintaining the structural arrangement of the headpiece that is characteristic of the E2 state. Nevertheless, the RMSD plot shows a 0.1-0.2 nm difference in the average RMSD between the lipid-free and lipid-bound states, which suggests that the lipid binding modulates the structural dynamics of the headpiece. Therefore, we plotted the interdomain distance distributions between residues Val223–Lys515 (A–N domains), Val223–Asp351 (A–P domains), and Lys515–Asp351 (N–P domains) to analyze the structural dynamics of the cytosolic headpiece more quantitatively. We found that in the presence of nonannular lipid, the interdomain distances are generally similar to those calculated in the crystal structure of the pump, indicating that the spatial arrangement of the A–N, A–P and N–P interfaces in the crystal structure of the E2 states is similar to the average geometry observed in solution (Figure 6C). In the absence of nonannular lipid, these interdomain distances increase by up to 0.4 nm with respect to the distances calculated from the crystal structure of the pump (Figure 6C). In particular, the distance histogram for N–A domains shows that these domains become dynamically more disordered in the absence of nonannular lipid (Figure 6A). We found that there is a substantial overlap in the distance values for the domain pairs A–P and N–P in the presence and absence of the nonannular lipid (Figure 6C), indicating that nonannular lipid removal has a marginal effect on the interdomain structural dynamics of A–P and N–P.

**Figure 6.**
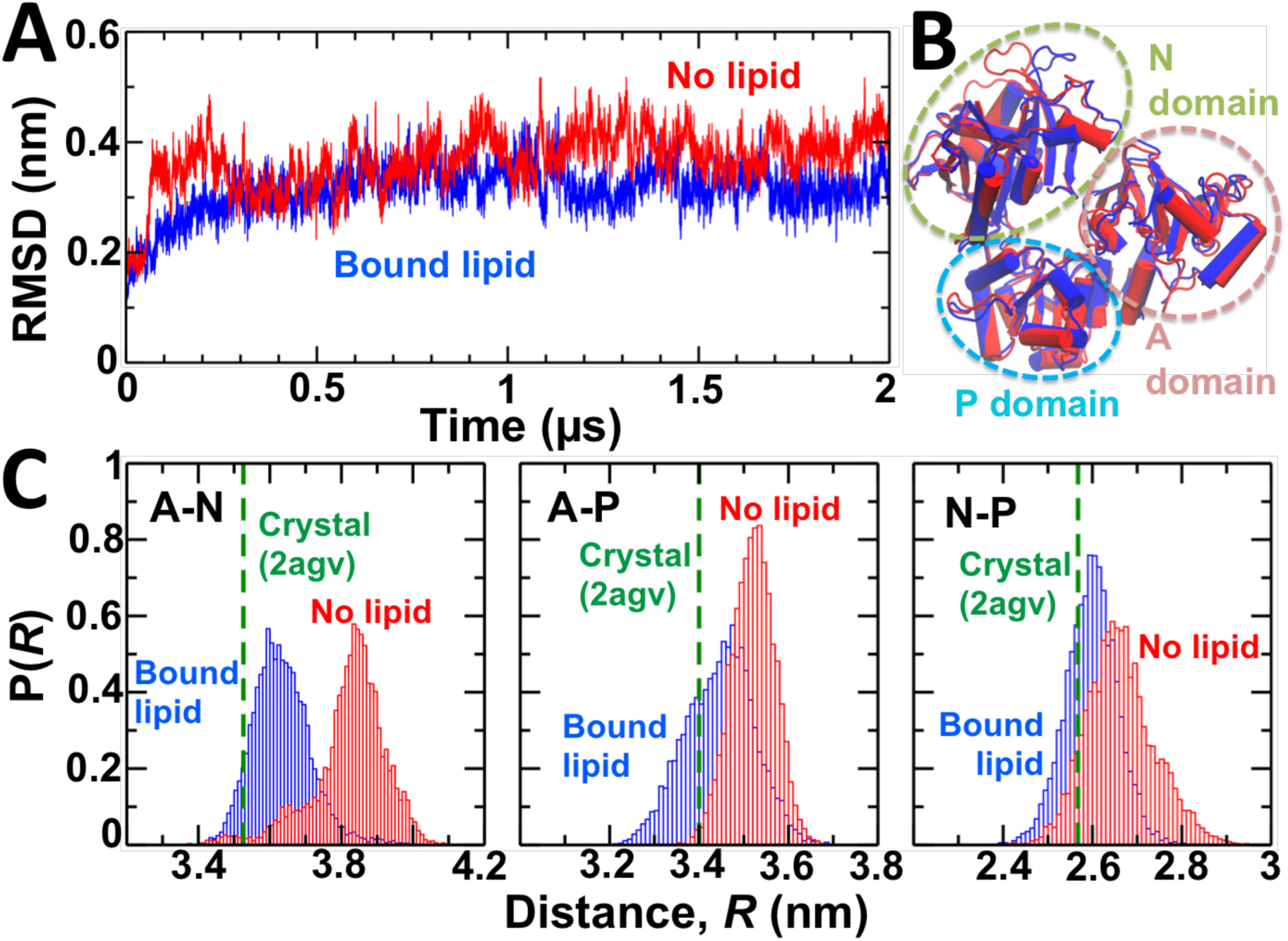
Structural dynamics of the cytosolic headpiece of SERCA in the presence and absence of nonannular lipid. (A) RMSD evolution of the headpiece of nonannular lipid-bound (blue) and nonannular lipid-free (red) SERCA. The RMSD of the headpiece was calculated by aligning the backbone of the cytosolic headpiece with the crystal structure of the pump (2agv^35^). (B) Superimposition of the most representative structures of the cytosolic headpiece of SERCA in the presence (blue) and absence (red) of nonannular lipid. The dashed ovals show the location of each functional domain of the cytosolic headpiece: N (green oval), P (blue oval), and A (brown oval). (C) Interdomain distance distributions between residues Val223–Lys515 (A–N domains), Val223–Asp351 (A–P domains), and Lys515–Asp351 (N–P domains). The dashed line shows the distance calculated from the crystal structure of the pump.

The changes in the distance distribution between the N and A domains suggest that nonannular lipid binding probably plays a role in the allosteric control of the intrinsic structural dynamics of the E2 state, but analysis of distance distributions alone is not sufficient to demonstrate this effect. Therefore, we used Cartesian principal component analysis (cPCA) to determine whether the shifts in the interdomain distance distributions induced by nonannular lipid removal actually reflect changes in the collective atomic motions of the pump. In both nonannular lipid-bound and nonannular lipid-free SERCA, first 10 and 35 principal components account for 90% and 95% of total atomic positional fluctuations, respectively (Figure 7A). Interestingly, we found remarkable similarity in the contribution of the principal components between lipid-bound and lipid-free SERCA. Probability distributions for the first 10 PCs extracted in the trajectories shows that both nonannular lipid-bound and lipid-free SERCA, the first two principal components belong to the essential space (*r*^2^<0.9), while the third principal component retains some non-Gaussian features (0.9<*r*^2^<0.98) (Figure 7B). In both trajectories, all consecutive principal components belong to the non-essential space (*r*^2^³0.98) (Figure 7B). The quantitative similarities between the two trajectories suggests that the conformational space of SERCA is not altered by the presence or absence of nonannular lipid. To further test this hypothesis, we calculated the inner products to quantify the mutual collinearity of individual principal components calculated from each MD trajectory. We found that first three principal components calculated from each independent MD trajectory are highly collinear (inner product values between 0.8 and 0.9), while lower collinearity is found further down in the PC hierarchy (Figure 7C). Despite these differences, the high collinearity of the three first principal components, which represent the essential phase space of lipid-bound and lipid-free SERCA, indicates that the nonannular lipid does not modulate the intrinsic structural dynamics of the E2 state of the pump.

**Figure 7.**
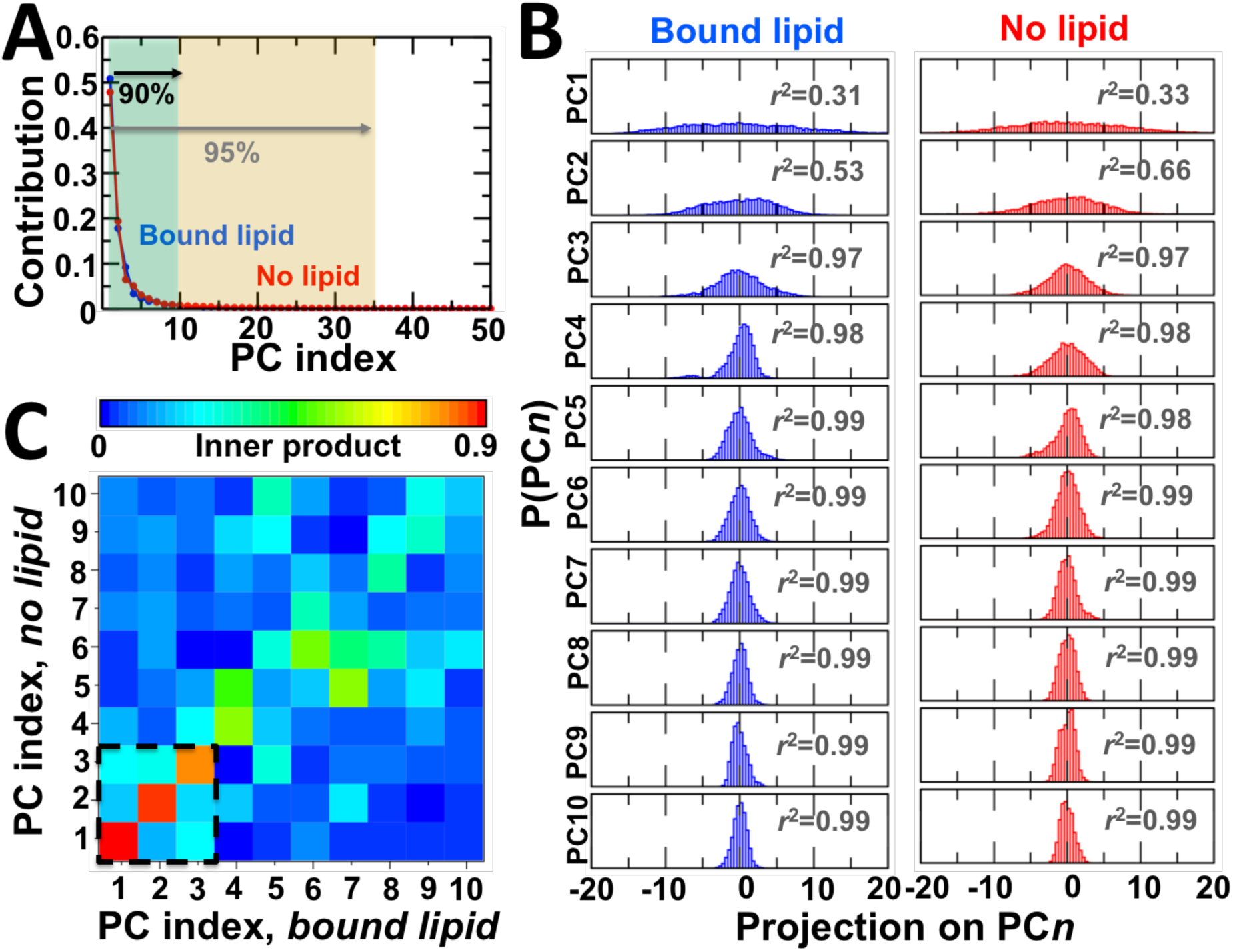
Effects of nonannular lipid binding on the intrinsic dynamics of the E2 state. (A) Contribution of the first 50 principal components to the variance in the structural space of SERCA in the presence (blue) and absence (red) of nonannular lipid. In both independent MD trajectories, we found that the first 10 and 35 principal components account nearly equally for 90% and 95% of total atomic positional fluctuations, respectively. (B) Probability distributions for the first 10 PCs extracted in the trajectories of nonannular lipid-bound (blue) and nonannular lipid-free (red) E2 state of SERCA. The *r*^2^ value shown inside each plot corresponds to the coefficient of determination obtained after fitting each histogram to a one-Gaussian distribution by least squares analysis. (C) Inner product matrix of the first ten principal component vectors of SERCA in the presence and absence of nonannular lipid. A value close or equal to 1 indicates that the principal components exist in the same dimension (e.g., structural sub-space); values close or equal to 0 indicate dissimilarity in the structural space captured by a given principal component.

### Effects of nonannular lipid binding on destabilization of the E2 state

Destabilization of the E2 state, which is required for producing a competent SERCA structure required for coupling ATPase activity and Ca^2+^ transport, initiates with proton release from residue Glu309 to the cytosol through transient water wires^41^. Removal of the nonannular lipid binding induces the formation of a more compact environment around Glu309 (Figure 4), so it is possible that nonannular lipid binding mediates the formation of these water wires, thus facilitating or inhibiting SERCA activation. Therefore, we searched for water wires between within a N-terminal pore that connects directly the sidechain of Glu309 with the cytosol via a narrow pathway formed around transmembrane helices M1, M2 and M4^41^.

Water wires are formed inside the N-terminal pore in the presence and absence of nonannular lipid (Figure 8). In general, water wires adopt similar structural arrangements, where three hydrogen-bonded water molecules occupy this pore to link the carboxyl group of Glu309 and the carboxamide group of Asn101. In the presence of nonannular lipid, we detected the formation of 6 water wire formation events, whereas 8 water wires are formed in the absence of the bound lipid. In both trajectories, we found that the water wire lifetime fluctuates between 100 and 150 ps, and that the lifetime is independent of lipid binding. The configuration, temporal scale and lifetime of these water wires is consistent with previous studies of the pump^41^, and indicate that the binding of the nonannular lipid has no effect on deprotonation and subsequent destabilization of the E2 state.

**Figure 8.**
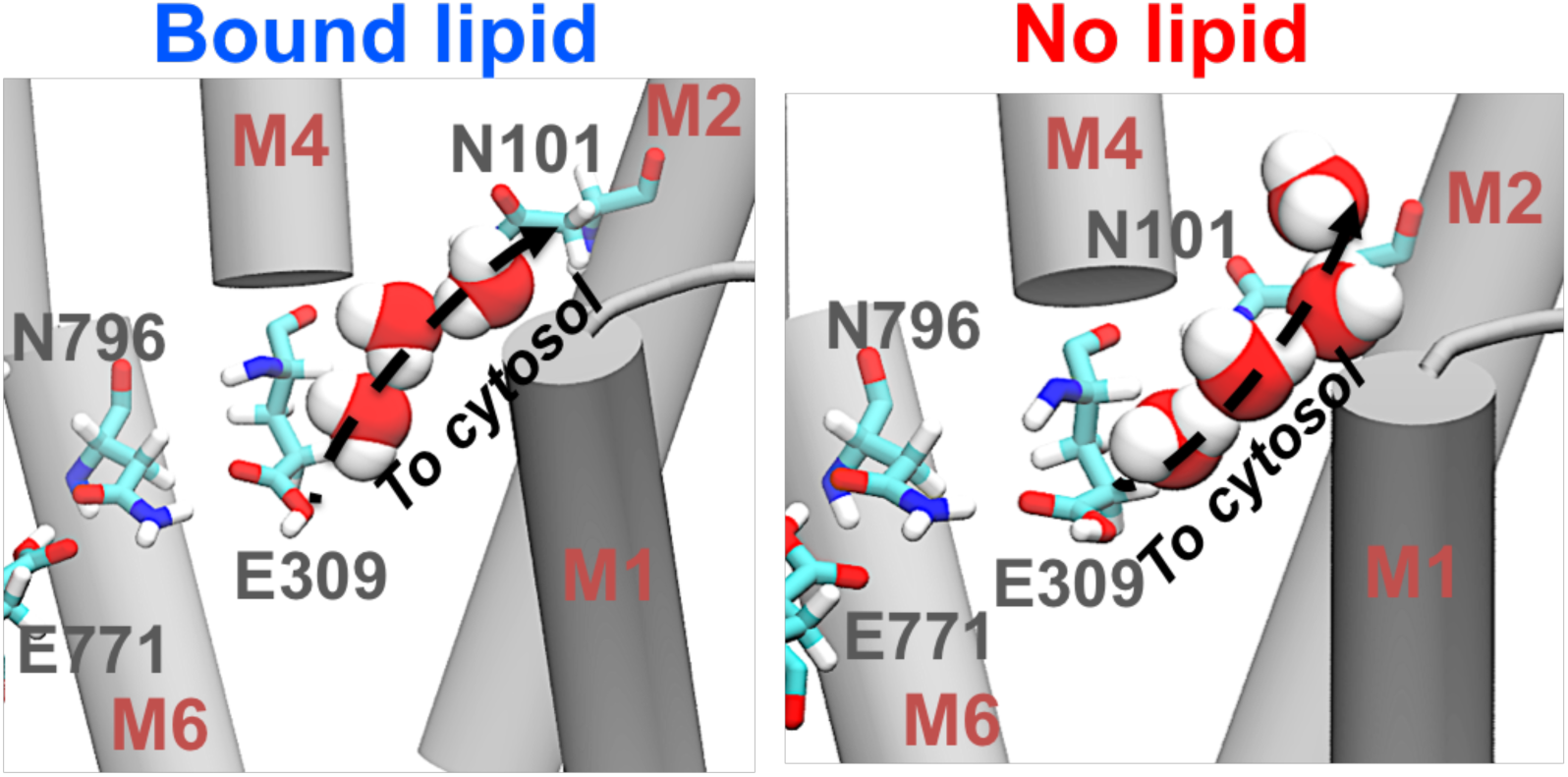
Structure of the water wires linking Glu309 with the cytosol in the presence and absence of nonannular lipid. In both cases, the hydrogen-bonded water wires link the transport site residue Glu309 and Asn101 at the cytosolic side of the transmembrane domain. The arrow indicates the likely direction of proton translocation. SERCA’s transmembrane helices are shown as gray cartoon, SERCA residues are shown as sticks, and water molecules are shown as spheres.

## Discussion

We report a mechanistic study of the effects of nonannular lipid binding on the stability of the E2 structural intermediate of SERCA. Previous studies have suggested that binding of a nonannular lipid acts as a wedge to keep M2 and M4 apart, thus stabilizing the E2 intermediate during the catalytic cycle of the pump^35^. Here, we characterized for the first time (to our knowledge) the effects of nonannular lipid binding on the stability of the E2 state of SERCA at physiological conditions and at an atomic level of detail. We show that in a physiological-like environment, the initially bound nonannular lipid remains bound to SERCA in the microsecond time scale, and its position is primarily stabilized by interactions between the lipid headgroup and polar residues of SERCA located in the vicinity of the phosphocholine group. Recent studies have shown that annular lipids form specific phospholipid–Arg/Lys and phospholipid–Trp interactions, and that these interactions have functional roles in the dynamics of SERCA^34^. However, we found that unlike the first layer of phospholipids that surround the transmembrane helices of the pump, the nonannular lipid does not establish a network of well-defined intermolecular interactions with SERCA, so it retains a substantial degree of mobility in solution. These findings are consistent with previous experimental studies suggesting that nonannular lipid binding to SERCA is transient and must be relatively unspecific to allow for efficient nonannular lipid dissociation during the E2-to-E1 transition of the pump^36^.

In the MD trajectories of the nonannular lipid-bound SERCA, the M2-M4 interface appears substantially more dynamic than that in the crystal structure of the pump. For example, the distance distribution between residues Gln108 (M2) and Thr316 (M4), indicates that the cytosolic regions of these transmembrane helices are in close contact during the trajectory, but also become more separated that in the crystal structure of the pump, e.g., *R*>1 nm in the simulation vs *R*=0.65 nm in the crystal structure. This finding is surprising because many of the nonannular protein–lipid interactions act as molecular glue that stabilizes protein–protein contacts, as in the V-type Na^+^ ATPase^43^. In the case of SERCA, we found that intramolecular protein–protein interactions between helices M2 and M4 are favored in the absence of nonannular lipid, and that lipid binding at this site disrupts these contacts. In the absence of nonannular lipid, the cytosolic segments of helices M2 and M4 are also in dynamic equilibrium between a spatially separated conformation and an arrangement in which both helices are in close physical contact. Comparison between the trajectories of lipid-bound and lipid-free SERCA further showed that lipid binding to the nonannular site does not have an effect on the helix-helix contacts along the luminal side of the transmembrane domain. These findings indicate that despite the mobility of the lipid in the nonannular site, binding of a lipid molecule induces physical separation of the cytosolic domains of transmembrane helices M2 and M4, in good qualitative agreement with previous studies showing that nonannular lipid binding induces a spatial separation of helices M2 and M4^36^.

We investigated whether binding of the nonannular lipid to SERCA stabilizes the E2 state by preventing the relative movement of transmembrane helices M2 and M4 of SERCA, but we found that occupancy of a lipid molecule in this site does not have any measurable effect on the structural dynamics of these regions of SERCA. Furthermore, the small structural rearrangement of the cytosolic M2–M4 interface detected in the absence of the nonannular lipid does not propagate to other transmembrane helices or the large cytosolic headpiece of the pump. These findings demonstrate that lipid binding at this site is not required for the structural stability of SERCA; instead, the structural stability of the E2 intermediate state is determined by protonation of the acidic residues in the transport sites, in agreement with previous studies^10,17,41,44^. While these findings indicate that nonannular lipid binding does not have a role in the stabilization of the E2 state, SERCA is susceptible to allosteric control by cations, small molecules, lipid composition and endogenous proteins, so it is possible that the nonannular lipid is an allosteric effector of the E2 state. Here, nonannular lipid binding may retard or accelerate activation of the E2-to-E1 transition through altering deprotonation of the E2 state^17,41^ or by inducing changes in the functional movements of the pump toward a competent E1 state^10^. Deprotonation of transport site residue Glu309 is key in destabilizing the E2 state, thus catalyzing the E2-to-E1 structural transition that is required for SERCA activation. Analysis of the MD simulations revealed that water wires connecting Glu309 and the cytosol are formed both in the presence and absence of nonannular lipid, and that there are no differences in the frequency and lifetime of these water wires. Recent studies by our group have shown that activation of the E2-to-E1 transition is largely determined by changes in the hierarchical organization and amplitude of rigid-body motions of the headpiece that ultimately shift the equilibrium between the E1 and E2 states. We found that the intrinsic structural dynamics of SERCA (e.g., the functional motions that determines the pump’s function^10,45^) are virtually identical in the presence and absence of the nonannular lipid. Together, these findings not only indicate that binding of a nonannular lipid molecule to SERCA is not required for the stability of the E2 intermediate state, but also suggest that occupancy of a lipid molecule at this site does not induce destabilization of this state. These findings are supported by extensive atomistic studies showing that activation of the E2-to-E1 transition occurs independently of nonannular lipid binding to SERCA^10^.

Finally, fluorescence quenching studies of SERCA reconstituted with a mixture of DOPC and brominated PC have suggested the alternative hypothesis that occupancy of nonannular lipids can lead to an increase in the activity of the pump^46^. Recent studies by our group showed that two lipid molecules are recruited by SERCA during the early structural transition between the E2 and the E1 states of the pump^16^. In this study, one lipid molecule binds rapidly (*t*<0.2 μs) to the nonannular lipid site described in this study; surprisingly, this lipid molecule adopts an orientation similar to that observed in crystal structures of the E2 state (**Supplementary Figure S1**) and remains bound to the pump in the microsecond timescale. The second nonannular lipid site is formed and occupied by a single lipid molecule at the site where the inhibitor thapsigargin binds to SERCA; this site is located near the cytosolic side of transmembrane helices M3 and M5 (**Supplementary Figure S1**). In both cases, these nonannular lipids appear to protect SERCA by ‘sealing’ transient hydrophobic patches in the transmembrane domain that result from the large structural changes in the pump during the E2-to-E1 transition^10^. Therefore, it is possible that binding of nonannular lipids actually facilitate SERCA activation through providing structural stability to the pump during the E2-to-E1 transition, which is a critical step required for the formation of a catalytically competent state of the pump^17^. We also do not rule out the possibility that nonannular lipids serve as structural building blocks that stabilize oligomeric states of the E2 state of the pump, since FRET experiments have shown that SERCA forms constitutive homodimers in living cells^47^. In this case, the large structural rearrangement in the transmembrane domain required for the E2-to-E1 transition probably disrupts protein sites required for oligomerization, so binding of a nonannular lipid might protect or stabilize these sites during the transition to prevent changes in the fraction of the SERCA oligomers and alterations in the conformational coupling of the pump^47^.

## Conclusions

We have used all-atom μs MD simulations to probe the effects of nonannular lipid binding on the stability of the calcium pump SERCA. We have tested the crystallography-based hypothesis that nonannular lipid binding is required for the structural stability of the E2 intermediate state of the pump. Our data indicates that the structural integrity of the E2 state is independent of nonannular lipid binding, and that protonation of the transport sites alone is sufficient to stabilize this intermediate. Moreover, extensive structural analysis suggests that occupancy of a lipid molecule at the nonannular site does not exert direct control on the destabilization of the E2 state required to initiate the E2-to-E1 transition. While these findings do not suggest a direct functional role of nonannular lipid binding to SERCA, we provide several alternative hypotheses that can be tested by experiments and simulations. For instance, it is possible that binding of a nonannular lipid can stabilize a structural intermediate along the E2-to-E1 transition, or that lipid binding is required to preserve protein–protein sites in the transmembrane domain that are required for oligomerization of the pump. Overall, we conclude that nonannular lipid binding is not necessary for the stability of the E2 state, but we speculate that it becomes functionally significant during the E2-to-E1 transition of the pump.

## Methods

### Modeling and setting up of the protonated E2 state of SERCA bound to a nonannular lipid

All crystal structures reported to date of SERCA1a bound to a nonannular lipid include the polar headgroup of the lipid only^34^, so we first modeled the structure of the E2 state bound to the full-length nonannular lipid. We docked a single POPC lipid molecule onto the crystal structure of the E2 state with the highest available resolution (PDB: 2agv^35^; resolution: 0.24 nm); the polar headgroup of the best docked orientation showed an excellent overlap with the position of the crystallographic lipid (RMSD=0.25 nm). The final SERCA-lipid complex was subjected to energy minimization in vacuum, which further decreased the difference in orientation between the crystallographic and docked position of the lipid headgroup to a RMSD value of 0.15 nm. This lipid bound structure of SERCA was used as an initial structure to simulate the effects of nonannular lipids on the stability and structural dynamics of the E2 state. On the basis of previous studies of the E2 state of the pump^40,41^, transport site residues Glu309, Glu771, and Glu908 were modeled as protonated, and residue Asp800 was modeled as ionized. The nonannular lipid-bound SERCA structure was inserted in a preequilibrated 12 × 12 nm POPC bilayer; we used the first layer phospholipids that surround SERCA in the E1 state^34^ as a reference to insert the complex in the lipid bilayer. This SERCA-lipid system was immersed in a rectangular box of TIP3P water, with a minimum margin of 1.5 nm between the protein and the edges of the periodic box in the *z*-axis. Finally, K^+^ and Cl^-^ ions were added to produce concentrations of 100 mM and 110 mM, respectively.

### Setting up E2 SERCA in the absence of nonannular lipid

We used 0.24-nm resolution crystal structure of the E2 state as a starting structure to simulate SERCA in the absence of nonannular lipid; the PDB codes of the structures used is 2avg^35^. We set up SERCA using similar chemical conditions and lipid composition used for nonannular lipid-bound SERCA to determine the effects of nonannular lipid removal on the stability and structural dynamics of the protonated E2 state. Here, the protonation states of transport sites residues were set up as follows: Glu309, Glu771 and Glu908 were modeled as protonated, whereas Asp800 was modeled as ionized. The SERCA-bilayer system was solvated using TIP3P water molecules, and K^+^ and Cl^-^ ions were added in the same concentrations used for the SERCA-nonannular lipid system.

### Molecular dynamics simulations

Atomistic simulations of the protonated E2 systems in the presence and absence of a nonannular lipid were performed by using the program NAMD^48^. For the simulations, we used with periodic boundary conditions^49^, particle mesh Ewald for calculating electrostatic interactions in periodic molecular systems^50,51^, a non-bonded cutoff of 0.9 nm, and the RATTLE algorithm^52^ to allow a time step of 2 fs. CHARMM36 force field topologies and parameters were used for the SERCA^53^, lipid^54^, water, K^+^, and Cl^-^. The NPT ensemble was maintained with a Langevin thermostat (310K) and an anisotropic Langevin piston barostat (1 atm). Fully solvated systems were first subjected to energy minimization, followed by gradually warming up of the systems for 200 ps. This procedure was followed by 10 ns of equilibration with harmonic restraints applied to the heavy atoms of the protein and nonannular lipid atoms (when applicable) using a force constant of 1000 kcal mol^-1^ nm^-2^. Finally, each SERCA system was simulated without restraints for a total simulation time of 4 μs.

### Analysis and visualization

We used the program VMD^55^ for structural analysis of the trajectories and visualization. We also analyzed the formation of water wires from residue Glu309 to the cytosol to determine whether nonannular lipid binding inhibits or facilitates proton release from the E2 state. Based on previous studies reporting the cytosolic proton release pathways in the E2 state, we measured the formation of two proton release pathways: a N-terminal pathway formed between Glu309 and Asn101^41^, and proton transfer from Glu309 to residues Asp800/Glu908, located at the beginning of a C-terminal pathway^56^. We define a transmembrane water wire as any configuration of hydrogen-bonded waters in a pore or channel connecting Glu309 with the cytosol. Water wire formation was determined as a wire/no-wire process irrespective of the actual number of hydrogen-bonded configurations or the identity of the water molecules.

### Principal component analysis

We used cPCA to characterize the essential space of SERCA, and to determine whether the structural changes observed in our MD simulations are associated with stabilization or destabilization of the E2 state. cPCA uses the actual dynamics of the protein to generate the appropriate collective coordinates that capture the important structural and dynamic features of the native states of the protein^57^. For cPCA, we first aligned SERCA structures using the 10-helix transmembrane domain as a reference. We then projected the trajectories of the SERCA into the phase space; the projection of a trajectory on the eigenvectors of its covariance matrix is called principal component. We then constructed probability histograms of each principal component; each histogram is then fitted to a one-Gaussian distribution to determine whether a principal component belongs to essential phase space^57^. Distributions with a coefficient of determination *r*^2^ <0.9 is considered to be non-Gaussian, and thus belong to the essential phase space^58^. Distributions with *r*^2^ values between 0.9 and 0.98 indicate that the principal component retains some significant non-Gaussian features, therefore contribute to the essential phase space. Finally, principal component distributions with *r*^2^>0.98 are Gaussian fluctuations, thus not considered part of the essential phase space^58^. All cPCA calculations and analyses were performed using the GROMACS package^59^.

## Acknowledgments

This work was supported by the National Institutes of Health grant R01GM120142. This work used the Extreme Science and Engineering Discovery Environment (XSEDE), which is supported by National Science Foundation Grant ACI-1548562.

## Data Availability

The datasets generated during and/or analyzed in the current study are available from the corresponding author on reasonable request.

